# Beyond comparisons of means: understanding changes in gene expression at the single-cell level

**DOI:** 10.1101/035949

**Authors:** Catalina A. Vallejos, Sylvia Richardson, John C. Marioni

## Abstract

Single-cell RNA sequencing (scRNA-seq) can be used to characterise differences in gene expression patterns between pre-specified populations of cells. Traditionally, differential expression tools are restricted to the study of changes in overall expression between cell populations. However, such analyses do not take full advantage of the rich information provided by scRNA-seq. In this article, we present a Bayesian hierarchical model which can be used to study changes in expression that lie beyond comparisons of means. In particular, our method can highlight genes that undergo changes in cell-to-cell heterogeneity between the populations but whose overall expression is preserved. Evidence supporting these changes is quantified using a probabilistic approach based on tail posterior probabilities, where a probability cut-off is calibrated through the expected false discovery rate. Our method incorporates a built-in normalisation strategy and quantifies technical artefacts by borrowing information from technical spike-in genes. Control experiments validate the performance of our approach. Finally, we compare expression patterns of mouse embryonic stem cells between different stages of the cell cycle, revealing substantial differences in cellular heterogeneity.

## Background

The transcriptomics’ revolution — moving from bulk samples to single-cell resolution — provides novel insights into a tissue’s function and regulation. In particular, single-cell RNA sequencing (scRNA-seq) has led to the identification of novel sub-populations of cells in multiple contexts [26, 11, 20]. However, compared to bulk RNA-seq, a critical aspect of scRNA-seq datasets is an increased cell-to-cell variability among the expression counts. Part of this variance inflation is related to biological differences in the expression profiles of the cells (e.g. changes in mRNA content and the existence of cell sub-populations or transient states), which disappears when measuring bulk gene expression as an average across thousands of cells. Nonetheless, this increase in variability is also due in part to *technical noise* arising from the manipulation of small amounts of starting material, which is reflected in weak correlations between technical replicates [4]. Such technical artefacts are confounded with genuine transcriptional heterogeneity and can mask biological signal.

Among others, one objective of RNA-seq experiments is to characterise transcriptional differences between pre-specified populations of cells (given by experimental conditions or cell-types). This is a key step for understanding a cell’s fate and functionality. In the context of bulk RNA-seq, two popular methods for this purpose are edgeR [23] and DESeq [1]. Nonetheless, these are not designed to capture features that are specific to scRNA-seq datasets. In contrast, SAMstrt [13] and SCDE [14] have been specifically developed to deal with scRNA-seq datasets. All of these methods target the detection of *differentially expressed genes* based on log fold changes (LFC) of overall expression between the populations. However, restricting the analysis to changes in overall expression does not take full advantage of the rich information provided by scRNA-seq. In particular — and unlike bulk RNA-seq — scRNA-seq can also reveal information regarding cell-to-cell expression heterogeneity. Critically, traditional approaches will fail to highlight genes whose expression is less stable in any given population but whose overall expression remains unchanged.

More flexible approaches, capable of studying changes that lie beyond comparisons of means are required in order to better characterise differences between distinct populations of cells. In this article, we develop a quantitative method to fill this gap, allowing the identification of genes whose cell-to-cell heterogeneity pattern changes between pre-specified populations of cells. This analysis allows the identification of genes with less variation in expression levels within a specific population of cells, which might indicate that they are under more stringent regulatory control. Additionally, genes with increased biological variability in a given population of cells could suggest the existence of additional sub-groups within the analysed populations. To the best of our knowledge, this is the first probabilistic tool developed for this purpose in the context of scRNA-seq analyses. We demonstrate the performance of our method using control experiments and by comparing expression patterns of mouse embryonic stem cells (mESCs) between different stages of the cell cycle.

## Results and discussion

### A statistical model to detect changes in expression patterns for scRNA-seq datasets

We propose a statistical approach for the comparison of expression patterns between *P* prespecified populations of cells. It builds upon BASiCS [25], a Bayesian model for the analysis of scRNA-seq data. As in traditional differential expression analyses, for any given gene i, changes in overall expression are identified by comparing population-specific expression rates *μ_ip_* (*p* = 1,…, *P*). However, the main focus of our approach is to assess differences in biological cell-to-cell heterogeneity between the populations. These are quantified through changes in population-specific biological *over-dispersion* parameters *δ_ip_* (*p =* 1,…, *P*), designed to capture residual variance inflation (after normalisation, technical noise removal and adjustment for overall expression), avoiding the well-known confounding relationship between mean and variance in count-based datasets [16]. Importantly, such changes cannot be uncovered by standard differential expression methods, which are restricted to changes in overall expression. Hence, our approach provides novel biological insights by highlighting genes that undergo changes in cell-to-cell heterogeneity between the populations despite the overall expression level being preserved.

To disentangle technical from biological effects, we exploit *spike-in* genes that are added to the lysis buffer and thence theoretically present at the same amount in every cell (e.g. the 92 ERCC molecules developed by the External RNA Control Consortium [12]). These provide an internal control or “gold standard”, to estimate the strength of technical variability and to aid normalisation. In particular, these control genes allow inference on cell-to-cell differences in mRNA content, providing additional information about the analysed populations of cells. These are quantified through changes between cell-specific normalising constants 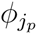 (the sub-index *j_p_* indicates the *j_p_*-th cell within the *p*-th population). A graphical representation of our model is displayed in Figure 1 (based on a two-groups comparison). It illustrates how our method borrows information across all cells and genes (biological transcripts and spike-in genes) in order to perform inference.

**Figure 1:**
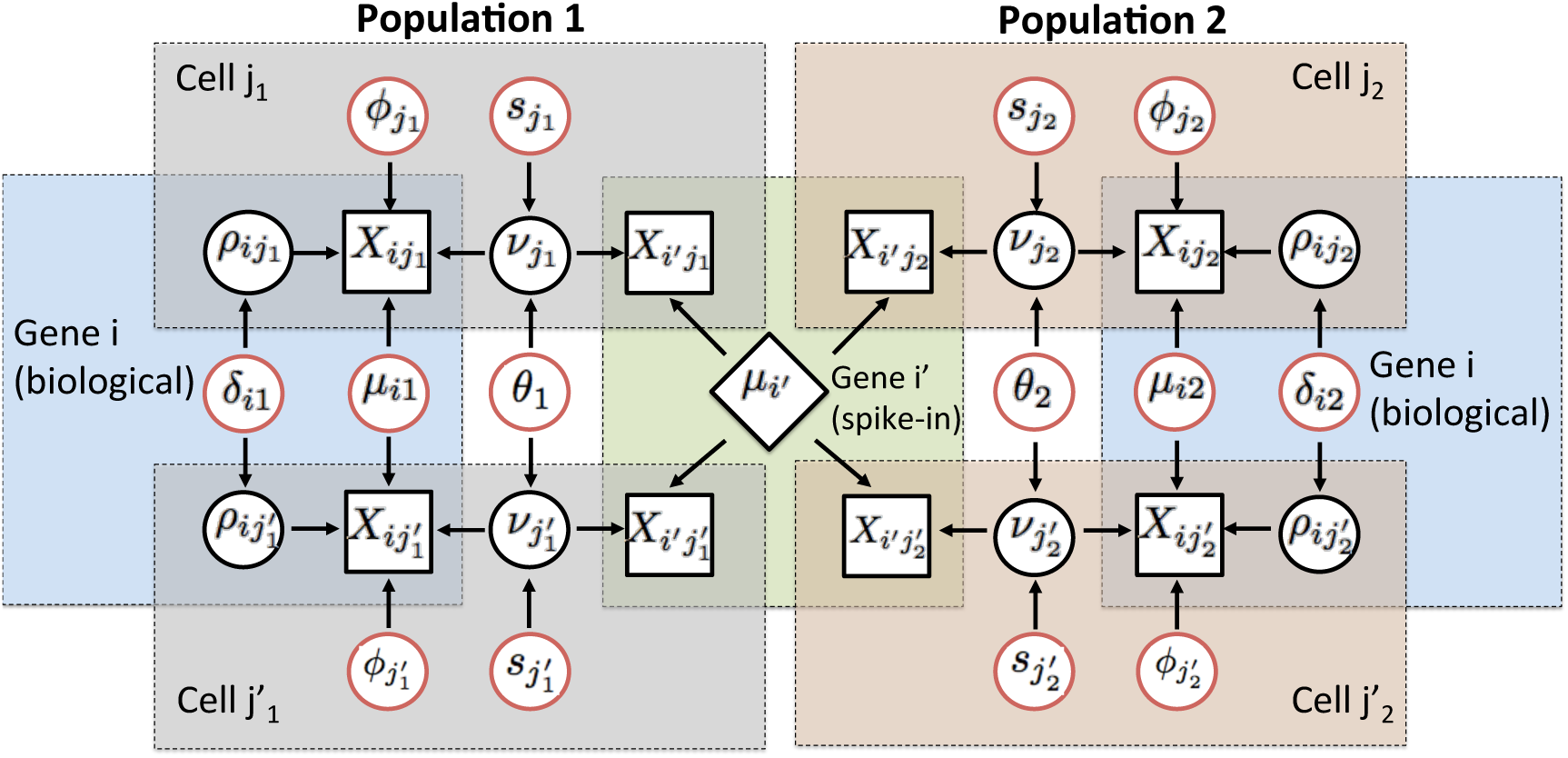
Graphical representation of our model for detecting changes in expression patterns (mean and over-dispersion) based on the comparison of two pre-defined population of cells. The diagram considers expression counts of 2 genes (*i:* biological and *i*′: technical) and 2 cells (*j_p_* and 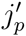) from each population *p* = 1, 2. Observed expression counts are represented by square nodes. The central rhomboid node denotes the known input number of mRNA molecules for a technical gene *i′*, which is assumed to be constant across all cells. The remaining circular nodes represent unknown elements, using black to denote random effects and red to denote model parameters (fixed effects) that lie on the top of the model’s hierarchy. Here, 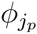 and 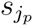 act as normalising constants that are cell-specific and 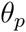 are global over-dispersion parameters capturing technical variability, which affects the expression counts of all genes and cells within each population. Finally, 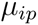 and 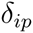 respectively measure overall expression of a gene *i* and its residual biological cell-to-cell over-dispersion (after normalisation, technical noise removal and adjustment for overall expression) within each population. Coloured areas highlight elements that are shared within a gene and/or cell. The latter emphasises how our model borrows information across all cells to estimate parameters that are gene-specific and all genes to estimate parameters that are cell-specific. More details regarding the model setup can be found in the Methods section of this article.

Posterior inference is implemented via a Markov Chain Monte Carlo (MCMC) algorithm, generating draws from the posterior distribution of all model parameters (see supplementary material). Post-processing of these draws allows quantification of supporting evidence regarding changes in expression patterns (mean and over-dispersion). These are measured using a probabilistic approach based on tail posterior probabilities, where a probability cut-off is calibrated through the expected false discovery rate (EFDR) [19]. Our strategy is flexible and can be combined with a variety of decision rules. As an illustration, the examples described in this article focus on the detection of genes whose absolute LFC in biological cell-to-cell heterogeneity between populations *p* and *p'* exceeds a minimum tolerance threshold *ω*_0_, i.e. when 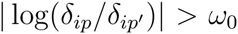 for a given *ω*_0_ ≥ 0. Similarly, our examples assess changes in overall expression using the decision rule 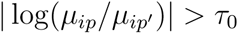, where *τ*_0_ ≥ 0 is an *a priori* chosen threshold for LFC in overall expression. These LFC thresholds can be used, e.g. to avoid highlighting genes with small changes in expression which are likely to be less biologically relevant.

More details regarding the model setup and the implementation of posterior inference can be found in the Methods.

### A control experiment: comparing single-cells versus pool-and-split samples

To demonstrate the efficacy of our method, we use the control experiment described in [9], where single mouse embryonic stem cells (mESCs) are compared against *pool-and-split* samples, consisting of pooled RNA from thousands of mESCs split into single-cell equivalent volumes. Such a controlled setting provides a situation where substantial changes in overall expression are not expected as, on average, the overall expression of single cells should match the levels measured on pool-and-split samples. Additionally, the design of pool-and-split samples removes biological variation, leading to a homogenous set of samples. Hence, pool-and-split samples are expected to show a genuine reduction in biological cell-to-cell heterogeneity when compared to single-cells.

Here, we display the analysis of samples cultured in a *2i* media. In these data, expression counts correspond to the number of molecules mapping to each gene within each cell. This is achieved by using unique molecular identifiers (UMIs), which remove amplification biases and reduce sources of technical variation [10]. Our analysis includes 74 single cells and 76 pool-and-split samples (same inclusion criteria as in [9]) and expression counts for 9,378 genes (9,343 biological and 35 ERCC spikes) defined as those with at least 50 detected molecules in total across all cells. As expected, our method does not reveal major changes in overall expression between single-cells and pool-and-split samples as the distribution of LFC estimates is roughly symmetric with respect to the origin (see upper panels in Figure 2) and the majority of genes are not classified as differentially expressed at 5% EFDR (see Figure 3b). However, this analysis suggests that setting the minimum LFC tolerance threshold *τ*_0_ equal to 0 is too liberal as small log-fold changes are associated with high posterior probabilities of changes in expression (see Figure 3a) and the number of differentially expressed genes is inflated (see Figure 3b). In fact, counter-intuitively, 4,710 genes (~ 50% of all analysed genes) are highlighted to have a change in overall expression when using *τ*_0_ = 0 (a similar issue is also observed when analysing bulk RNA-seq data [17]). In contrast, this number is reduced to 559 genes (~ 6% of all analysed genes) when setting *τ*_0_ = 0.4.

**Figure 2:**
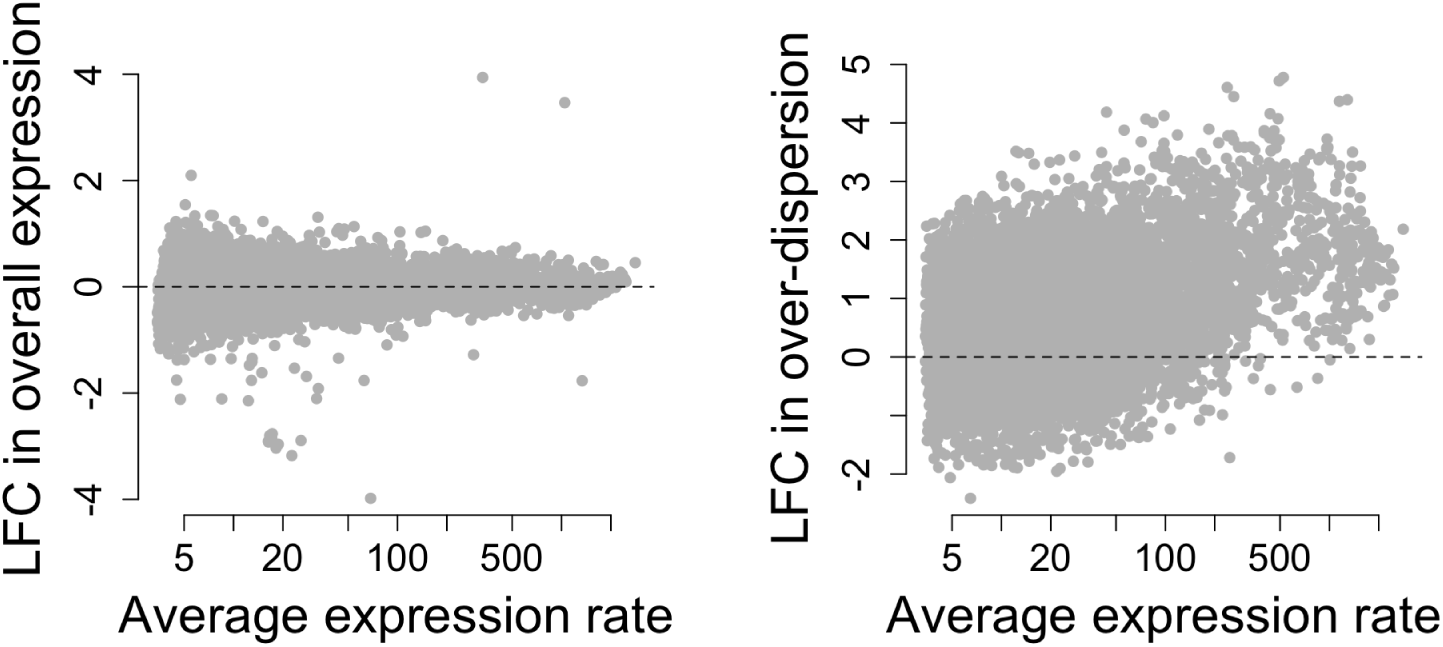
Estimated LFCs in expression (mean and over-dispersion) when comparing single-cells versus pool-and-split samples (*2i* serum culture). Posterior medians of LFC in overall expression log(*μ_i_*_SC_/*μ_i_*_P&s_) (left panel) and biological over-dispersion log(*μ_i_*_SC_/*μ_i_*_P&s_)(right panel) against the average between estimates of overall expression rates for single-cells (SC) and pool-and-split (P&S) samples. Average values are defined as a weighted average between groups, with weights given by the number of samples within each group of cells. As expected, our analysis does not reveal major changes in expression levels between SC and P&S samples. In fact, the distribution of estimated LFCs in overall expression is roughly symmetric with respect to the origin. In contrast, we infer a substantial decrease in biological over-dispersion in the P&S samples. This is reflected by a skewed distribution of estimated LFCs in biological overdispersion towards positive values.

**Figure 3:**
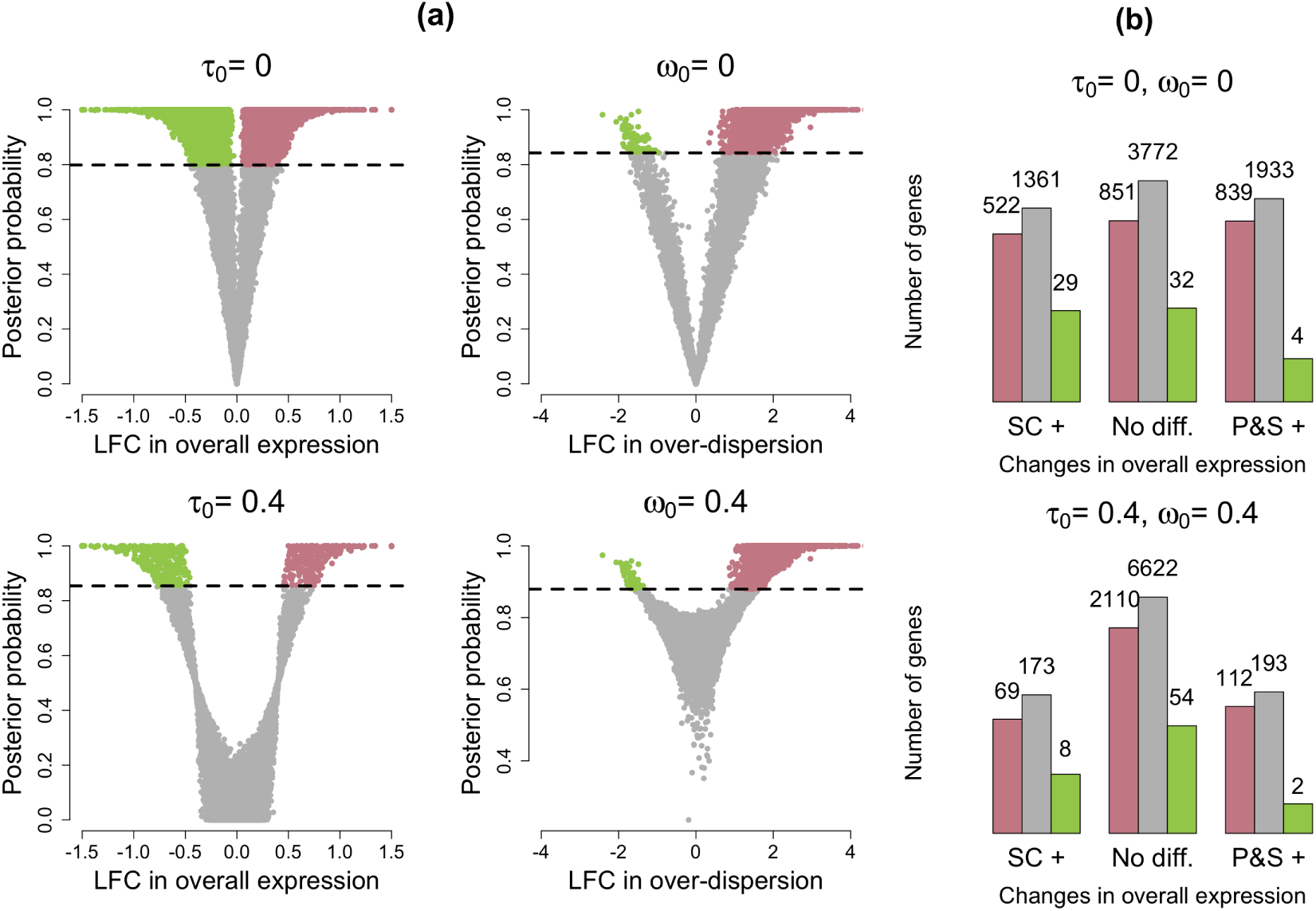
Summary of changes in expression patterns (mean and over-dispersion) for single-cells versus pool-and-split samples (EFDR = 5%). (a) Volcano plots showing posterior medians of LFCs against estimated tail posterior probabilities. Left panels relate to the test where we assess if the absolute LFC in overall expression between single-cells (SC) and pool-and-split (P&S) samples exceeds a minimum threshold *τ*_0_. Estimates for LFCs in overall expression are truncated to the range (–1.5,1.5). Pink and green dots represent genes highlighted to have higher overall expression in the SC and P&S samples, respectively. Right panels relate to the test where we assess if the absolute LFC in biological over-dispersion between SC and P&S samples exceeds a minimum threshold *ω*_0_. In all cases, horizontal dashed lines are located at probability cutoffs defined by EFDR = 5%. Pink and green dots represent genes highlighted to have higher biological over-dispersion in the SC and P&S samples, respectively. (b) Bins in the horizontal axis summarise changes in overall expression between the groups. We use SC+ and P&S+ to denote higher overall expression was detected in SC and P&S+ samples, respectively (the central group of bars (No diff.) corresponds to those genes where no significant differences were found). Coloured bars within each group summarise changes in biological over-dispersion between the groups. We use pink and green bars to denote higher biological over-dispersion in SC and P&S+ samples, respectively (and grey to denote no significant differences were found). Number of genes are displayed in log-scale.

Posterior inference regarding biological over-dispersion is consistent with the experimental design, where the pool-and-split samples are expected to have more homogeneous expression patterns. In fact, as shown in the right hand panel of Figure 2, the distribution of estimated LFCs in biological over-dispersion is skewed towards positive values (higher biological over-dispersion in single-cells). This is also supported by the results shown in Figure 3b, where slightly more than 2,000 genes exhibit increased biological over-dispersion in single cells and almost no genes (~ 60 genes) are highlighted to have higher biological over-dispersion in the pool-and-split samples (EFDR = 5%). In this case, the choice of *ω*_0_ is less critical (within the range explored here). This is illustrated by the left panels in Figure 3a, where tail posterior probabilities exceeding the cut-off defined by EFDR = 5% correspond to similar ranges of LFC estimates.

### mESCs across different cell cycle stages

Our second example shows the analysis of the mESC dataset presented in [5], which contains cells for which the cell cycle phase is known (G1, S and G2M). After applying the same quality control criteria as in [5], our analysis considers 182 cells (59, 58 and 65 cells in stages G1, S and G2M, respectively). To remove genes with consistently low expression across all cells, we excluded those genes with less than 20 RPM, on average, across all cells. After this filter, 5,687 genes remain (including 5,634 intrinsic transcripts, and 53 ERCC spike-in genes). As a proof of concept, to demonstrate the efficacy of our approach under a negative control, we performed permutation experiments, where cell labels were randomly permuted into 3 groups (containing 60, 60 and 62 samples, respectively). In such a case, our method correctly infers that mRNA content as well as gene expression profiles do not vary across groups of randomly permuted cells (see Figure 4).

**Figure 4:**
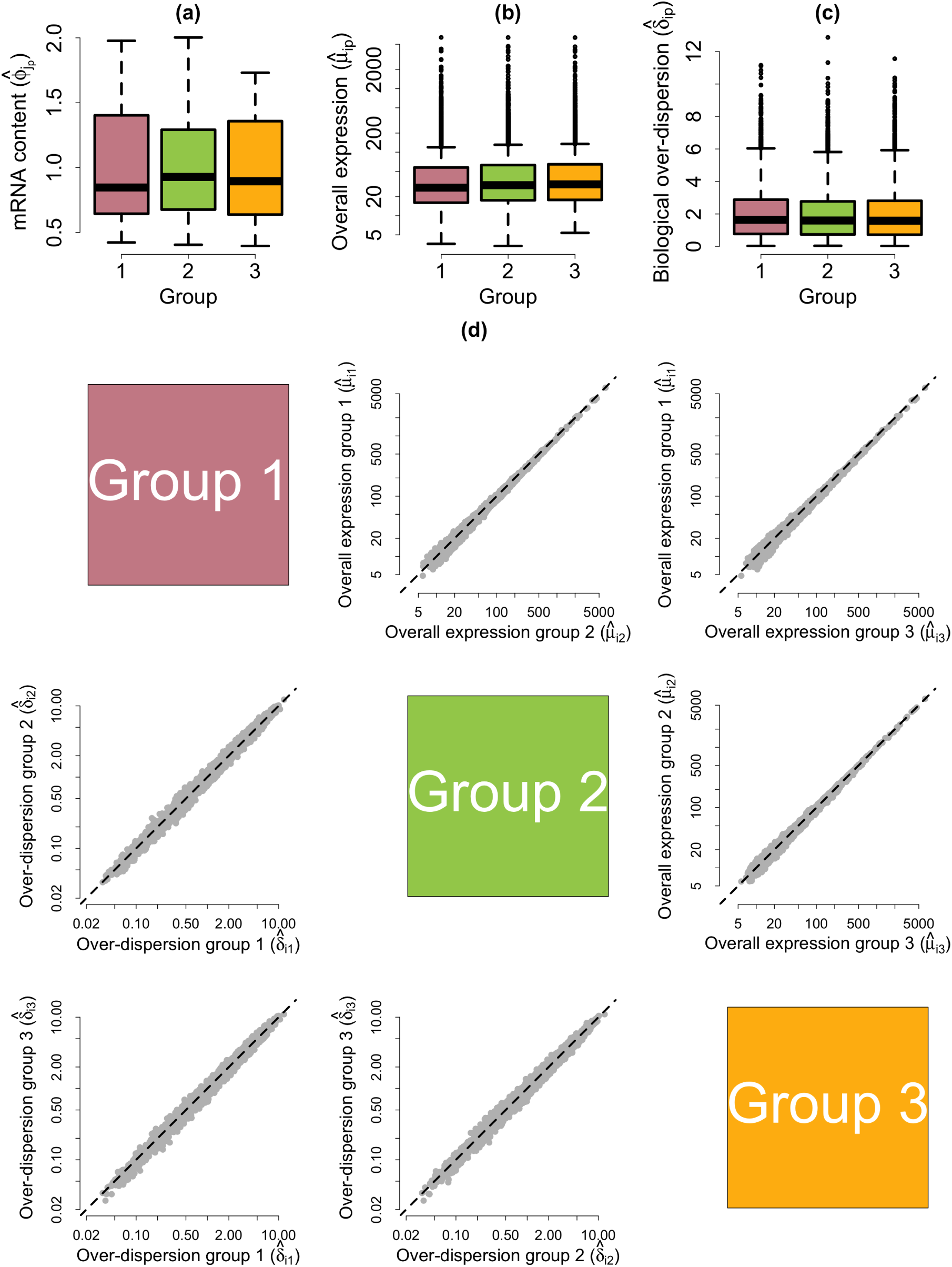
Posterior estimates of model parameters based on random permutations of the mESC cell cycle dataset. For a single permuted dataset (a) Empirical distribution of posterior medians for mRNA content normalising constants 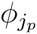· across all cells. (b) Empirical distribution of posterior medians for gene-specific expression rates 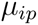across all genes. (c) Empirical distribution of posterior medians for gene-specific biological over-dispersion parameters *δ_ip_* across all genes. (d) As an average across 10 random permutations. Upper diagonal panels compare estimates for gene-specific expression rates *μ_ip_* between groups of cells. Lower diagonal panels compare gene-specific biological over-dispersion parameters *δ_ip_* between groups of cells.

As cells progress through the cell cycle, cellular mRNA content increases. In particular, our model infers that mRNA content is roughly doubled when comparing cells in G1 versus G2M, which is consistent with the duplication of genetic material prior to cell division (see Figure 5a). After normalisation, we rule out a global shift in expression levels between cell cycle stages (see Figure 5b and upper triangular panels in Figure 5d). Nonetheless, a small number of genes are identified as displaying changes in overall expression between cell cycle phases at 5% EFDR (see Figure 6). The actual number of differentially expressed genes depends on whether we use *τ*_0_ = 0 or *τ*_0_ = 0.4. However, the permutations described above suggest that *τ*_0_ = 0.4 is more appropriate as, just by chance, it is likely to observe LFCs below this threshold (in fact, *τ*_0_ = 0.4 roughly coincides with the 90-th percentile of the empirical distribution of posterior estimates for LFC in overall expression across all genes and permutations). Additionally, this LFC threshold is equivalent to a minimum 50% increase in overall expression (in whichever group the gene has higher expression). To validate our results, we performed Gene Ontology (GO) enrichment analysis within those genes classified as differentially expressed between cell cycle phases (see supplementary material). Here, we discuss results related to *τ*_0_ = 0.4. Not surprisingly, we found an enrichment of mitotic genes among the 545 genes classified as differentially expressed between G1 and G2M cells. In addition, the 209 differentially expressed genes between S and G2M are enriched for regulators of cytokinesis, which is the final stage of the cell cycle where a progenitor cell is divided into two daughter cells [7].

**Figure 5:**
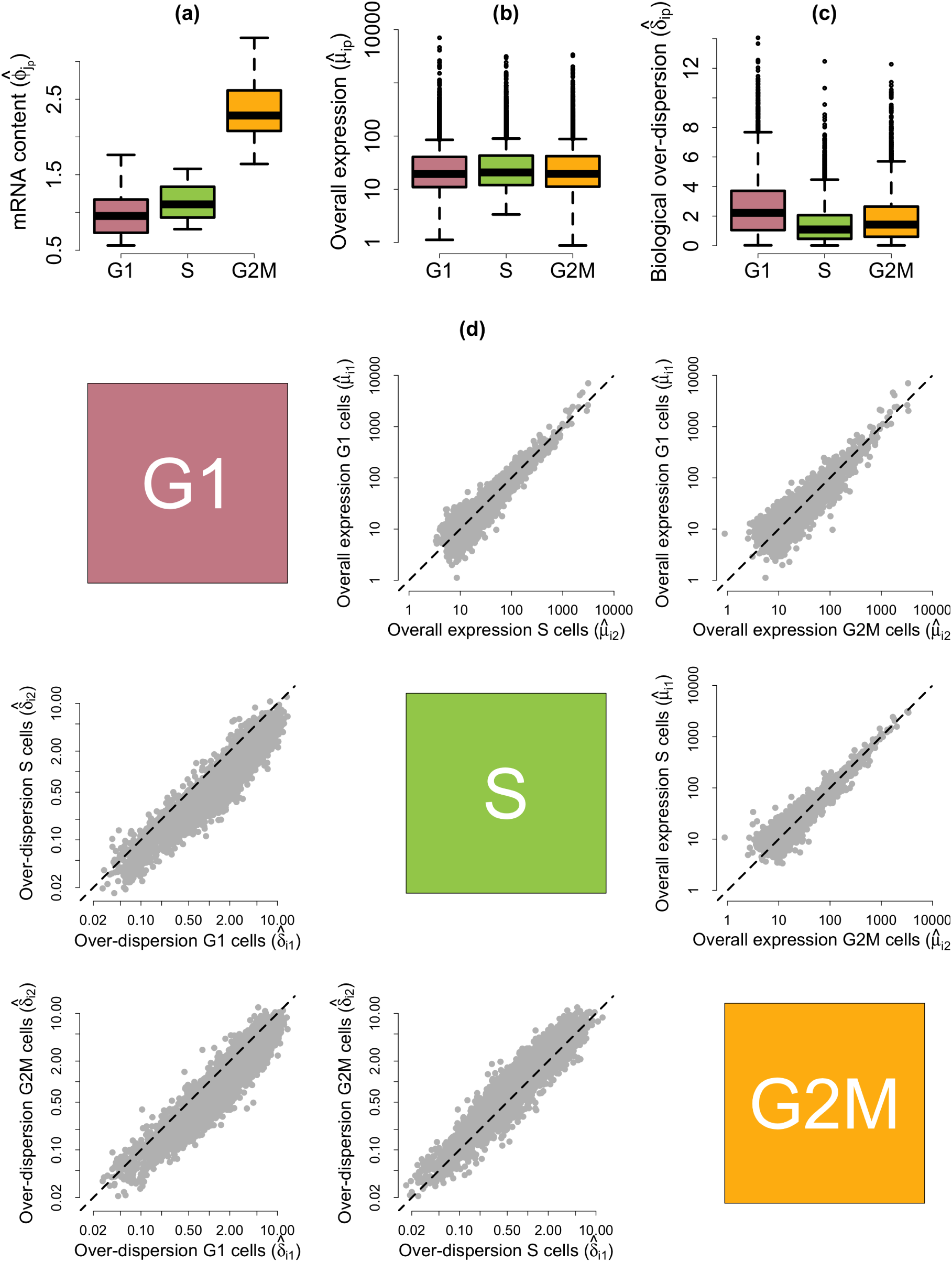
Posterior estimates of model parameters for mESCs across different cell cycle phases. (a) Empirical distribution of posterior medians for mRNA content normalising constants 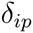 across all cells. (b) Empirical distribution of posterior medians for gene-specific expression rates *μ_ip_* across all genes. (c) Empirical distribution of posterior medians for gene-specific biological over-dispersion parameters *δ_ip_* across all genes. (d) Upper diagonal panels compare estimates for gene-specific expression rates *μ_ip_* between groups of cells. Lower diagonal panels compare gene-specific biological over-dispersion parameters *δ_ip_* between groups of cells.

**Figure 6:**
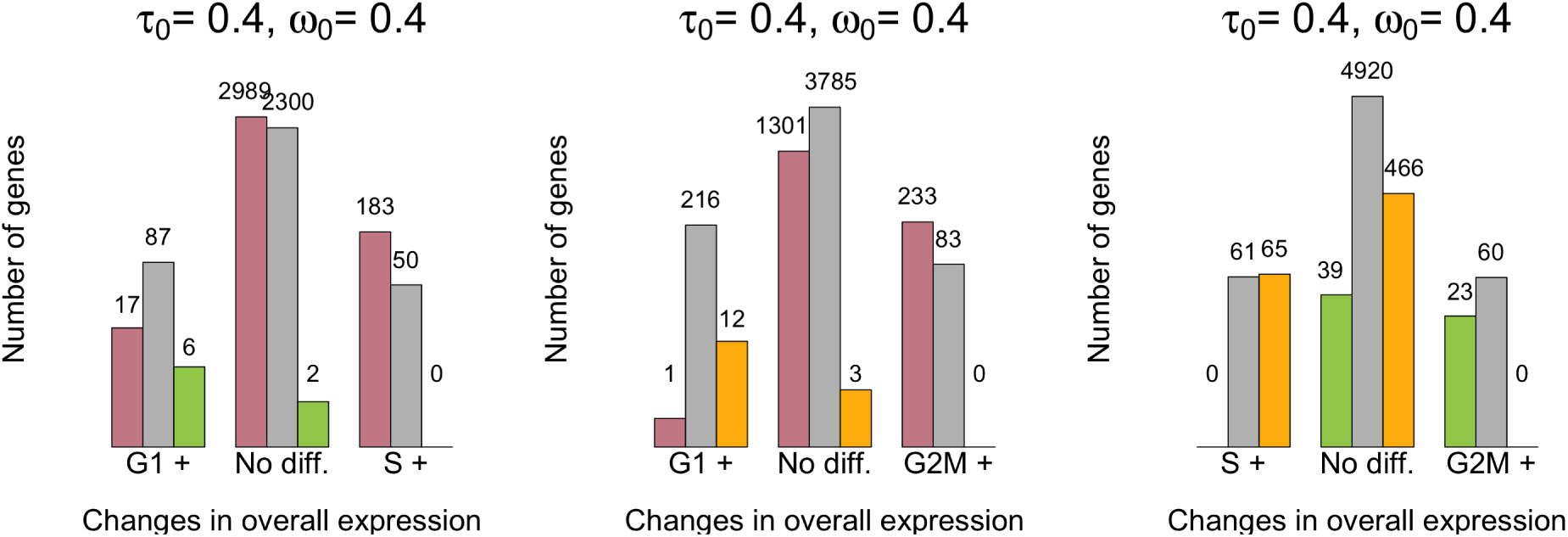
Summary of changes in expression patterns (mean and over-dispersion) for the mESC cell cycle dataset (EFDR = 5%). Bins in the horizontal axis summarise changes in overall expression between each pair of groups. We use G1+, S+ and G2M+ to denote higher overall expression was detected in cell cycle phase G1, S and G2M, respectively (the central group of bars (No diff.) corresponds to those genes where no significant differences were found). Coloured bars within each group summarise changes in biological over-dispersion between the groups. We use pink, green and yellow bars to denote higher biological over-dispersion in cell cycle phases G1, S and G2M, respectively (and grey to denote no significant differences were found). Number of genes are displayed in log-scale.

Our method uncovers a substantial decrease in biological over-dispersion when cells move from G1 to S phase, followed by a slight increase after the transition from S to G2M phase (see Figure 5c and the lower triangular panels in Figure 5d). This is consistent with the findings in [7], where the increased gene expression variability observed in G2M cells is attributed to an unequal distribution of genetic material during cytokinesis and the S phase is shown to have the most stable expression patterns within the cell cycle. Here, we discuss GO enrichment of those genes whose overall expression rate remains constant but that exhibit changes in biological overdispersion between cell cycle stages (EFDR = 5%, *ω*_0_ = 0.4). Critically, these genes will not be highlighted by traditional differential expression tools, which are restricted to differences in overall expression rates. For example, among the genes with higher biological over-dispersion in G1 with respect to S phase, we found an enrichment of genes related to protein dephosphorylation. These are known regulators of the cell cycle [6]. Moreover, we found that genes with lower biological over-dispersion in G2M cells are enriched for genes related to DNA replication checkpoint regulation (which delays entry into mitosis until DNA synthesis is completed [3]) relative to G1 cells and mitotic cytokinesis when comparing to S cells. Both of these processes are likely to be more tightly regulated in G2M phase.

A full table with GO enrichment analysis of the results described here is provided in the supplementary material.

## Conclusions

Our method provides a quantitative tool to study changes in gene expression patterns between pre-specified populations of cells. Unlike traditional differential expression analyses, our model is able to identify changes in expression that are not necessarily reflected by shifts in mean. This allows a better understanding of the differences underlying distinct populations of cells. In particular, we focus on the detection of genes whose residual biological heterogeneity (after normalisation and technical noise removal) varies between the populations. This is quantified through biological over-dispersion parameters, which capture variance inflation with respect to the level that would be expected in a homogeneous population of cells. A decision rule is defined through a probabilistic approach based on tail posterior probabilities and calibrated using the expected false discovery rate. The performance of our method was demonstrated using a controlled experiment where we recovered the expected behaviour of gene expression patterns.

Currently, our approach requires pre-defined populations of cells (e.g. defined by cell types or experimental conditions). However, a large number of scRNA-seq experiments involve a mixed population of cells, where cell-types are not known *a priori* (e.g. [26, 11, 20]). In such cases, expression profiles can be used to *cluster* cells into distinct groups and to characterise markers for such sub-populations. Nonetheless, unknown group structures introduce additional challenges for normalisation and quantification of technical variability. A future extension of our work is to combine the estimation procedure within our model with a clustering step, propagating the uncertainty associated with each of these steps into downstream analysis.

Until recently, most scRNA-seq datasets consisted of hundreds (and sometimes thousands) of cells. However, droplet-based approaches [15, 18] have recently allowed parallel sequencing of substantially larger numbers of cells in an effective manner. This brings additional challenges to the statistical analysis of scRNA-seq datasets. In particular, current protocols do not allow the addition of technical spike-in genes. As a result, the deconvolution of biological and technical artefacts becomes less straightforward. Moreover, the increased sample sizes emphasise the need for more computationally efficient approaches that are still able to capture the complex structure embedded within scRNA-seq datasets. To this end, we foresee the use of parallel programming as a tool for reducing computing times. Additionally, we are also exploring approximated posterior inference based, for example, on an Integrated Nested Laplace Approximation (INLA) [24].

Finally, our approach lies within a generalised linear mixed model framework. Hence, it can be easily extended to include additional information such as covariates (e.g. cell cycle stage, gene length) and experimental design (e.g. batch effects) using fixed and/or random effects.

## Methods

### A statistical model to detect changes in expression patterns for scRNA-seq datasets

In this article, we introduce a statistical model for identifying genes whose expression patterns change between pre-defined populations (given by experimental conditions or cell types) of cells. Such changes can be reflected via the overall expression level of each gene as well as through changes in cell-to-cell biological heterogeneity. Our method is motivated by features that are specific to scRNA-seq datasets. In this context, it is essential to appropriately normalise and remove technical artefacts from the data before extracting biological signal. This is particularly critical when there are substantial differences in cellular mRNA content, amplification biases and other sources of technical variation. For this purpose, we exploit technical spike-in genes which are added at the same quantity to each cell’s lysate. A typical example is the set of 92 ERCC molecules developed by the External RNA Control Consortium [12]. Our method builds upon BASiCS [25] and can perform comparisons between multiple populations of cells using a single model. Importantly, our strategy avoids stepwise procedures where datasets are normalised prior to any downstream analysis. This is an advantage over methods using pre-normalised counts, as the normalisation step can be distorted by technical artefacts.

We assume that there are *P* groups of cells to be compared, each containing *n_p_* cells (*p* = 1…,*P*). Let 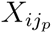 be a random variable representing the expression count of a gene *i* (*i* = 1…, *q*) in the *j_p_*-th cell from group p. Without loss of generality, we assume the first *q*_0_ genes are biological and the remaining *q* − *q*_0_ are technical spikes. Extending the formulation in BASiCS, we assume that

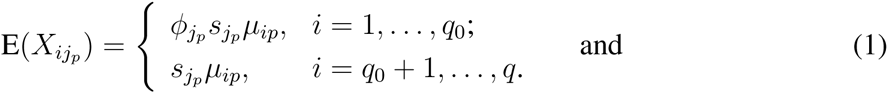

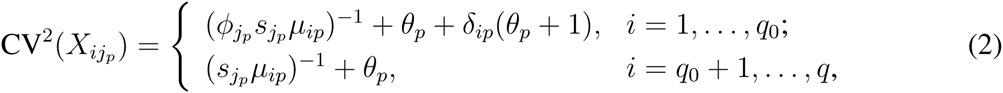

where CV stands for *coefficient of variation* (i.e. the ratio between standard deviation and mean). These expressions are the result of a Poisson hierarchical structure (see supplementary material). Here, 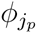 act as cell-specific normalising constants (fixed effects), capturing differences in input mRNA content across cells (reflected by the expression counts of intrinsic transcripts only). A second set of normalising constants, 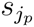, capture cell-specific scale differences affecting the expression counts of all genes (intrinsic and technical). Among others, these differences can relate to sequencing depth, capture efficiency and amplification biases. However, a precise interpretation of the 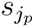 varies across experimental protocols, e.g. amplification biases are removed when using UMIs[10]. In addition, *θ_p_*’s are global technical noise parameters controlling the over-dispersion (with respect to Poisson sampling) of all genes within group *p*. The overall expression rate of a gene *i* in group *p* is denoted by *μ_ip_*. These are used to quantify changes in the overall expression of a gene across groups. Similarly, the *δ_ip_*’s capture residual over-dispersion (beyond what is due to technical artefacts) of every gene within each group. These so-called biological over-dispersion parameters relate to heterogeneous expression of a gene across cells. For each group, stable “*housekeeping-like*” genes lead to *δ_ip_ ≈* 0 (low residual variance in expression across cells) and highly variable genes are linked to large values of *δ_ip_*. A novelty of our approach is the use of *δ_ip_*’s to quantify changes in biological over-dispersion. Importantly — and unlike the CV — this avoids confounding effects due to changes in overall expression between the groups.

A graphical representation of this model is displayed in Figure 1. To ensure identifiability of all model parameters, we assume that *μ_ip_*’s are known in the case of the spike-in genes (and given by the number of spike-in molecules that are added to each well). Additionally, we impose the identifiability restriction

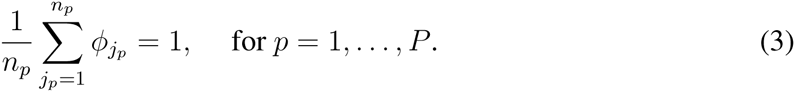

Here, we discuss the priors assigned to parameters that are gene and group-specific (see Additional file 2 for the remaining elements of the prior). These are given by

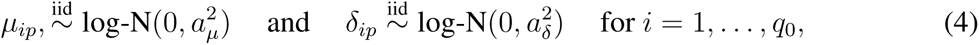

Hereafter, without loss of generality, we simplify our notation to focus on two-group comparisons. This is equivalent to assigning Gaussian prior distributions for LFCs in overall expression (r_i_) or biological over-dispersion (*ω_i_*). In such a case, it follows that

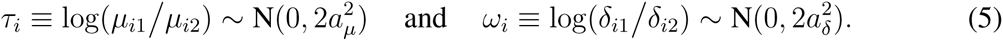

Hence, our prior is *symmetric*, meaning that we do not *a priori* expect changes in expression to be skewed towards either group of cells. Values for 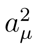 and 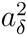 can be elicited using an *expected* range of values for LFC in expression and biological over-dispersion, respectively. The latter is particularly useful in situations where a gene is not expressed (or very lowly expressed) in one of the groups, where e.g. LFCs in overall expression are undefined (the maximum likelihood estimate of *τ_i_* would be ±∞, the sign depending on which group expresses gene *i*). A popular solution to this issue is the addition of *pseudo-counts*, where an arbitrary number is added to all expression counts (in all genes and cells). While the latter guarantees that *τ_i_* is well-defined, it leads to artificial estimates for *τ_i_* (see Table 1). Instead, our approach exploits an informative prior (indexed by 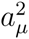) to *shrink* extreme estimates of *τ_i_* towards an expected range. This strategy leads to a meaningful shrinkage strength, which is based on prior knowledge. Importantly — and unlike the addition of pseudo-counts — our approach is also helpful when comparing biological over-dispersion between the groups. In fact, if a gene *i* is not expressed in one of the groups, this will lead to a non-finite estimate of *ω_i_* (if all expression counts in a group are equal to zero, the corresponding estimate of the biological over-dispersion parameters would be equal to zero). Adding pseudo-counts cannot resolve this issue, but imposing an informative prior for *ω_i_* (indexed by 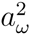) will shrink estimates towards the appropriate range.

Generally, posterior estimates of *τ_i_* and *ω_i_* are robust to the choice of 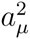 and 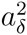, as the data is informative and dominates posterior inference. In fact, these values are only influential when shrinkage is needed, e.g. when there are zero total counts in one of the groups. In such cases, posterior estimates of *τ_i_* and *ω_i_* are dominated by the prior, yet the method described below still provides a tool to quantify evidence of changes in expression. As a default option, we use 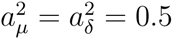.

Posterior samples for all model parameters are generated via an Adaptive Metropolis within Gibbs Sampling algorithm [22]. A detailed description of our implementation can be found in the supplementary material.

### Post-hoc correction of global shifts in input mRNA content between the groups

The identifiability restriction in (3) does only apply to cells within each group. As a consequence, if they exist, global shifts in cellular mRNA content between groups (e.g. if all mRNAs where present at twice the level in one population related to another) are absorbed by the *μ_ip_*’s. To correct this bias, we adopt a 2-step strategy where: (i) model parameters are estimated using the identifiability restriction in (3) and (ii) global shifts in input mRNA content are treated as a fixed *offset* and corrected post-hoc. For this purpose, we use the sum of overall expression rates (intrinsic genes only) as a proxy for the overall mRNA content within each group. Without loss of generality, we use the first group of cells as a reference population. For each population *p* (*p* = 1,…, *P*), we define a population specific offset effect

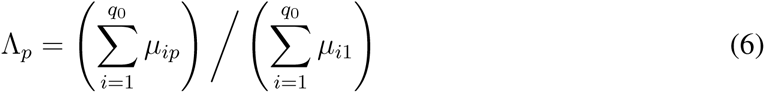

and perform the following offset correction

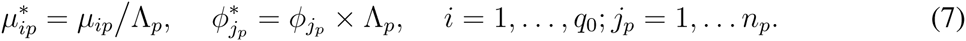

This is equivalent to replacing the identifiability restriction in (3) by

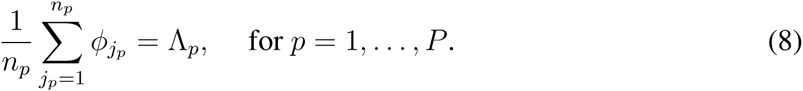

Technical details regarding the implantation of this post-hoc offset correction are explained in the supplementary material. The effect of this correction is illustrated in Figure 7 using the cell cycle dataset described in the main text. As an alternative, we also explored the use of the ratio between the total intrinsic counts over total spike-in counts to define a similar offset correction based on

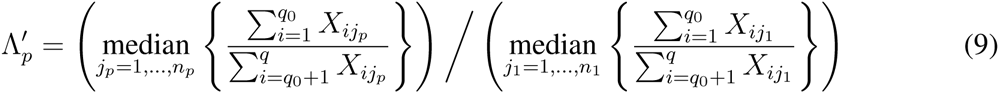

**Figure 7:**
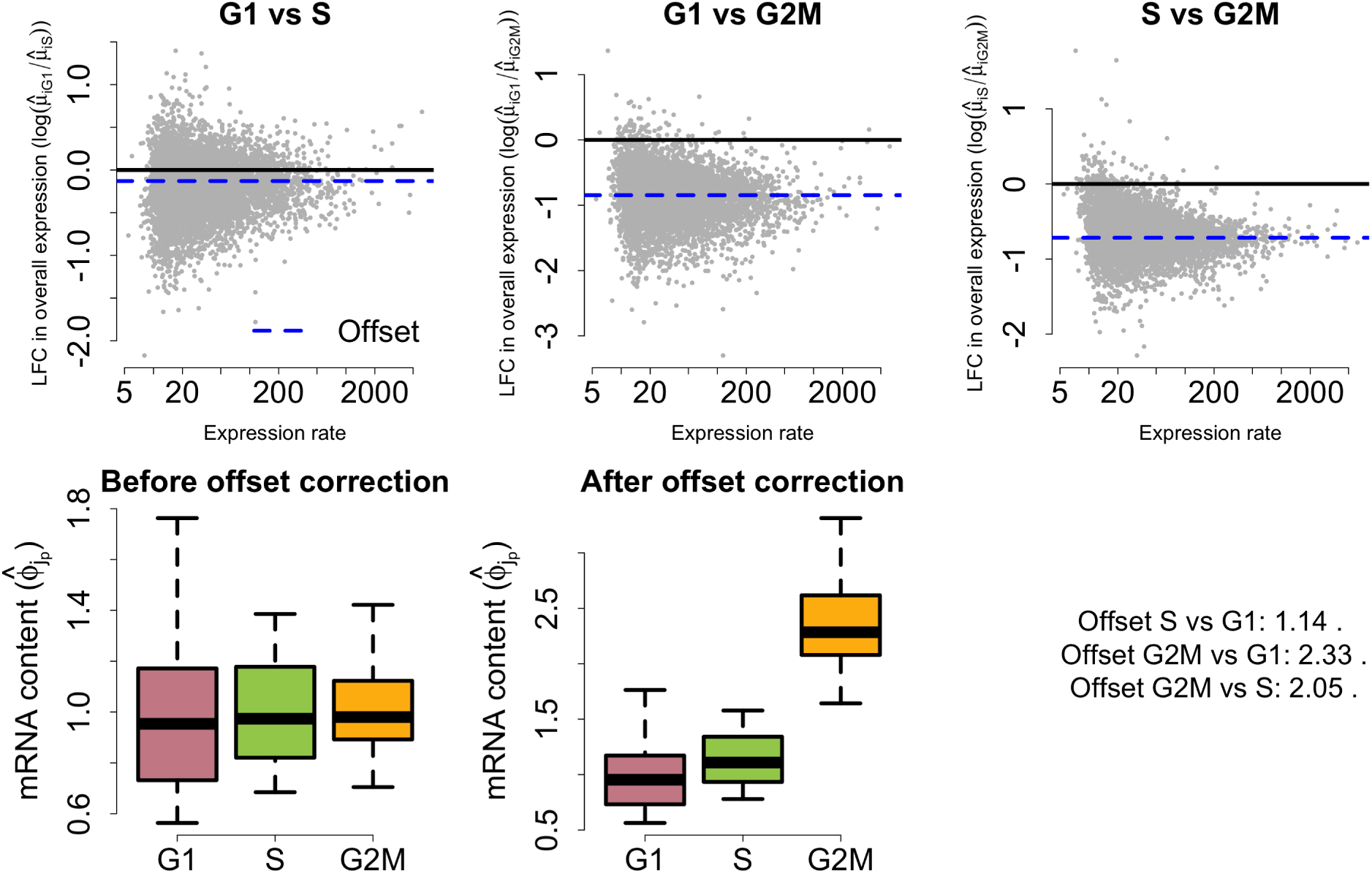
Post-hoc offset correction for cell cycle dataset. Upper panels display posterior medians for LFC in overall expression against the weighted average between estimates of overall expression rates for G1, S and G2M cells (weights defined by the number of cells in each group). Lower panels illustrate the effect of the offset correction upon the empirical distribution of posterior estimates for mRNA content normalising constants 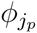. These figures illustrate a shift in mRNA content throughout cell cycle phases. In particular, our model infers that cellular mRNA is roughly duplicated when comparing G1 to G2M cells.

**Table 1:** Synthetic example to illustrate the effect of pseudo-counts addition over the estimation of log-fold changes in overall expression. For simplicity, we assume that normalisation is not required so that pseudo-counts are linearly reflected in the overall expression rates. While pseudo-counts introduce an additive effect, log-fold change estimates measure changes on a multiplicative scale. Hence, pseudo-counts addition leads to an artificial deflation of log-fold change estimates. As a consequence, such estimates cannot be meaningfully interpreted.

|  | Empirical<br>estimate | Adding 0.5<br>pseudo-counts | Adding 1<br>pseudo-count |
| --- | --- | --- | --- |
| Overall expression rate in population 1 ($\mu_{i1}$) | 10 | 10.5 | 11 |
| Overall expression rate in population 2 ($\mu_{i2}$) | 0 | 0.5 | 1 |
| Log-fold change in overall expression 1 vs 2 | $+\infty$ | 3.04 | 2.40 |

In the case of the cell cycle dataset, both alternatives are equivalent. Nonetheless, the first option is more robust in cases where a large number of differentially expressed genes are present. Hereafter, we use *μ_ip_* and 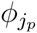 to denote 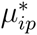 and 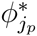, respectively.

### A probabilistic approach to quantify evidence of changes in expression patterns

A probabilistic approach is adopted, assessing changes in expression patterns (mean and overdispersion) through a simple and intuitive scale of evidence. Our strategy is flexible and can be combined with a variety of decision rules. In particular, here we focus on highlighting genes whose absolute LFC in overall expression and biological over-dispersion between the populations exceeds minimum tolerance thresholds *τ*_0_ and *ω*_0_, respectively (*τ*_0_, *ω*_0_ ≥ 0), set *a priori*.

For a given probability threshold *α_M_* (0.5 < *α_M_* < 1), a gene *i* is identified as exhibiting a change in overall expression between populations *p* and *p*′ if

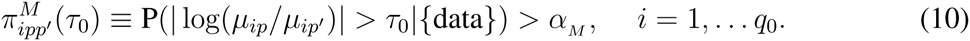

If *τ*_0_ → 0, 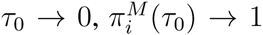 becoming uninformative to detect changes in expression. As in [2], in the limiting case where *τ*_0_ = 0, we define

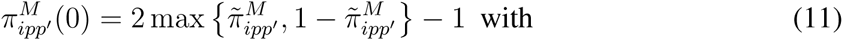

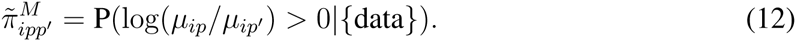

A similar approach is adopted to study changes in biological over-dispersion between populations *p* and *p*′, using

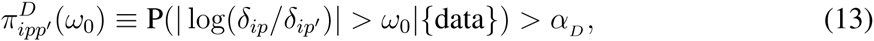

for a fixed probability threshold *α_D_* (0.5 < *α_D_* < 1). In line with (11) and (12), we also define

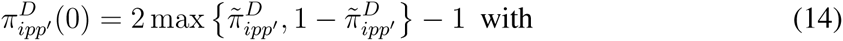

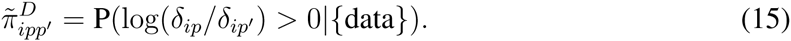

Evidence thresholds *α_M_* and *α_D_* can be fixed *a priori*. Otherwise, these can be defined by controlling the expected false discovery rate (EFDR) [19]. In our context, these are given by

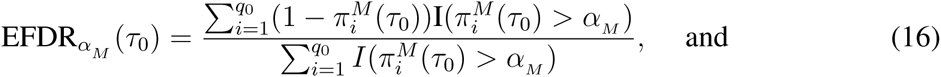

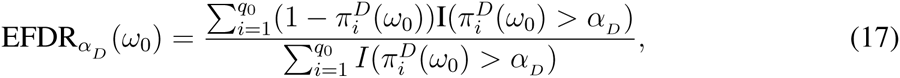

where I(*A*) = 1 if event *A* is true, 0 otherwise.

The posterior probabilities in (10), (11), (13) and (15) can be easily estimated — as a postprocessing step — once the model has been fitted (see supplementary material). In addition, our strategy is flexible and can be easily extended to investigate of more complex hypotheses, which can be defined post-hoc, e.g. to identify those genes that show significant changes in cell-to-cell biological over-dispersion but that maintain a constant level of overall expression between the groups.

### Software

Our implementation is freely available as an R package [21], using a combination of R and C++ functions through the Rcpp library [8].

This can be found in https://github.com/catavallejos/BASiCS.

## Availability of supporting data

All datasets analysed in this article are publicly available in the cited references.

## Author’s contributions

CAV, JCM and SR conceived and designed the methods and experiments. CAV implemented the methods and analysed the data. All authors were involved in writing the paper and have approved the final version.

## Acknowledgements

We acknowledge all members of Richardson research group (MRC-BSU) and Marioni laboratory (EMBL-EBI) for support and discussions during the preparation of this document. In particular, we are grateful to Nils Eling (EMBL-EBI), Antonio Scialdone (EMBL-EBI) and Aaron Lun (CRUK) for numerous discussions and suggestions than enriched the final version of the manuscript. JCM and CAV acknowledge core EMBL funding. SR and CAV acknowledge core MRC funding. JCM acknowledges core support from CRUK.

## Additional Files

Additional file 1 — Supplementary material

This document contain additional details regarding the statistical model presented in this article and the implementation of Bayesian inference. Additionally, it displays the R code used for the data analysis.

